# Association between land use, land cover, plant genera, and pollinator abundance in mixed-use landscapes

**DOI:** 10.1101/2021.01.20.427393

**Authors:** Vishesh L. Diengdoh, Barry W. Brook, Mark Hunt, Stefania Ondei

## Abstract

Pollinators are globally threatened by land-use change, but its effect varies depending on the taxa and the intensity of habitat degradation. However, pollinator-landscape studies typically focus on regions of intensive human activities and on a few focal species. Evaluating pollinator responses in landscapes with moderate land-use changes and on multiple pollinator groups would therefore fill an important knowledge gap. This study aims to determine the predictive capacity and effect of habitat characteristics on the relative abundance of multiple pollinator groups in mixed-use landscapes. To do this, we collected field data on the relative abundance of nectivorous birds, bees, beetles, and butterflies across the Tasman Peninsula (Tasmania, Australia). We then applied Random Forests to resolve the effects of land use (protected areas, plantation, and pasture), land cover at different radii (100 m and 2000 m), and plant genera on pollinator abundance. Overall, land cover and plant genera were more important predictors of pollinator abundance than land use. And the effect of land use, land cover, and plant genera varied depending on the pollinating group. Pollinator groups were associated with a range of plant genera, with the native genera *Acacia, Leptospermum, Leucopogon, Melaleuca, Pomaderris*, and *Pultenaea* being among the most important predictors. Our results highlight that one size does not fit all—that is pollinator response to different landscape characteristics vary, emphasise the importance of considering multiple habitat factors to manage and support a dynamic pollinator community, and demonstrates how land management can be informed using predictive modelling.

## Introduction

Pollinators are declining globally (Maes and Van Dyck 2001; Ollerton et al. 2014; Regan et al. 2015) due to the interactions and synergies between land-use change (habitat loss and fragmentation), introduced and invasive species, agrochemicals, and climate change (Potts et al. 2010; Regan et al. 2015; Vanbergen and Insect Pollinators Initiative 2013). Of these, land-use change is widely considered the most important threat to pollinators as it affects plant abundance and diversity, in turn reducing the availability of floral resources (Baude et al. 2016; Nicolson and Wright 2017; Paton 2000). This could have major cascading consequences for many habitats, as pollinators (animal vectors of pollen) are important ecosystem components and are estimated to globally pollinate 70% of crops (Klein et al. 2007) and 87% of wild plants (Ollerton et al. 2011).

While land-use change has an overall negative effect on pollinator abundance and richness (Winfree et al. 2011), this effect varies between different pollinating taxa, extent of landscape changes, and ecosystems (Millard et al. 2021; Montero-Castaño and Vilà 2012; Winfree et al. 2009). Yet, the literature on impact of landscape changes on pollinators is mostly focused on Hymenoptera (Senapathi et al. 2016) and in landscapes with extreme changes (Winfree et al. 2011). This raises the question of how bees and non-bee pollinating taxa would be impacted by landscapes subject to moderate alteration. Identifying the differences in the impacts of land-use types on different taxa would allow us to design better management policies.

Irrespective of the type of land use, plant species richness can have a positive impact on insect pollinators (Kral-O’Brien et al. 2021). For example, agricultural landscapes that include high-quality habitats support a higher abundance and richness of pollinators than agricultural landscapes without such habitats (Kavanagh et al. 2007; Kennedy et al. 2013). In this context, targeted plantings can be used to attract and sustain pollinators in degraded ecosystems (Menz et al. 2011). Pollinator restoration can be accomplished by using a subset of all available plant families or species (Campbell et al. 2019; Sabatino et al. 2021). That raises the question of whether there are plants (or plant groups) that are relatively more effective in sustaining pollinators and, if so, which species/characteristics are best suited for which taxa. Identifying and managing such plants would be beneficial in sustaining pollinator communities.

In this study, we aim to assess the predictive capacity and effect of land use, land cover, and plant genera on the relative abundance (count) of nectivorous birds, bees, beetles, and butterflies. To do this, we collected field data across protected areas, plantations, and pastures using the Tasman Peninsula (Tasmania, southern-temperate Australia) as a case study of a diverse, mixed-use landscape. We used Random Forests (Liaw and Wiener 2002) for predictive modelling, as it is a non-parametric decision-tree-based method commonly used in predictive analysis of complex, conditional data (Greenwell 2017). The advantage of using machine learning and predictive modelling is its ability to handle data with non-normal distribution and noise, deal implicitly with interactions, and use robust training-and-testing analysis to make predictions for informed decision making (Thessen 2016; Willcock et al. 2018). We discuss the associations between predictors and pollinators and which predictors should be managed to improve the diversity and resilience of pollinator communities.

## Methods

### Study area

The Tasman Peninsula, located in the south-eastern portion of the large island of Tasmania, Australia (Fig. 1), covers an area of 660.4 km^2^ with elevation from 0 to 582 m a.s.l. (meters above sea level). It is characterised by a mix of dry and wet sclerophyll eucalypt forest and dry coastal vegetation, and supports a third of the vascular plants found in Tasmania with 556 vascular plants, of which there are 336 dicotyledons (Brown and Duncan 1986). It was selected as a study area as its landscape experienced only moderate anthropogenic change relative to surrounding regions, since over a quarter (26.7%) of its area is protected (Australian Bureau of Statistics 2021). Overall, the study area is dominated by grazing pasture, forest plantations and protected areas (Department of Primary Industries Parks Water and Environment 2015) and lacks landscapes subject to extreme change, which are defined as areas with ≤5 % of native vegetation (Winfree et al. 2009).

**Fig. 1.**
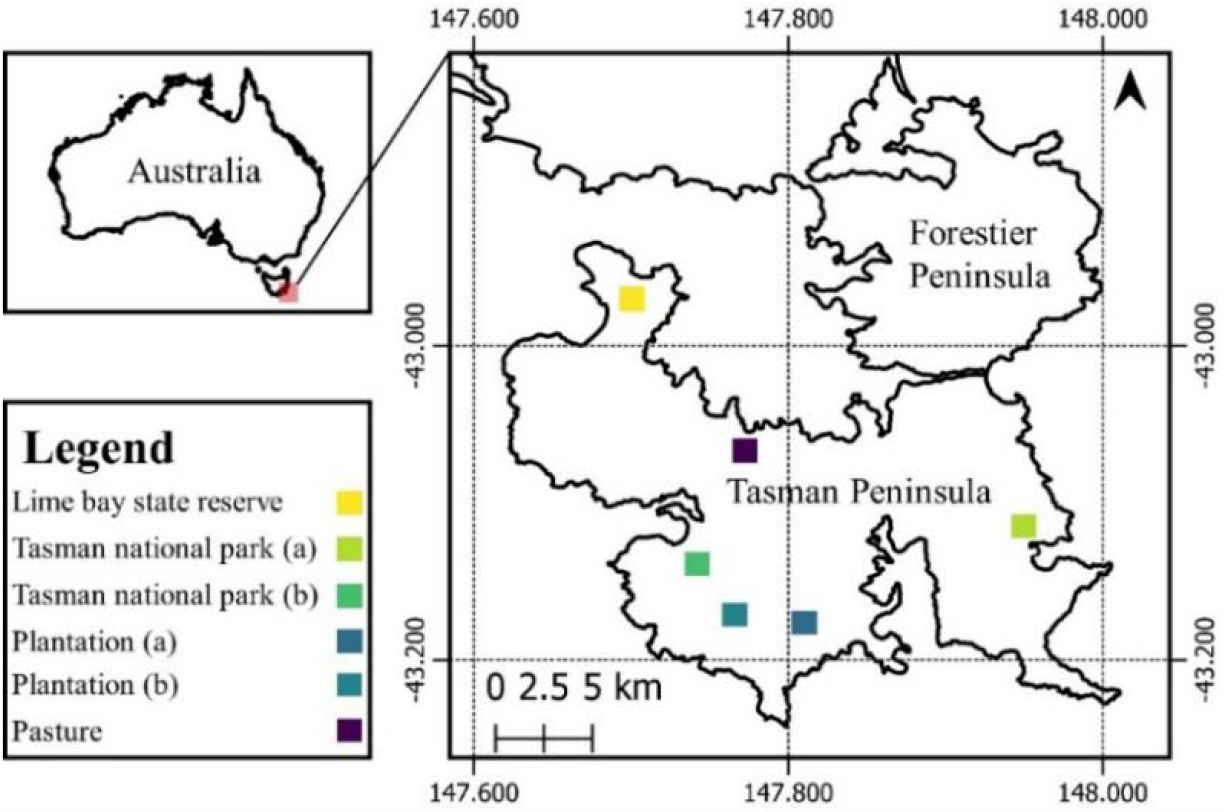
Location of the sampling land-use sites in the Tasman Peninsula, Tasmania, Australia.

### Study design

We established a total of 36 × 2-ha plots for bird observations. Of those, 18 plots were located in protected areas (six in each of the three protected areas), 12 in plantations (six each in of the two-plantations) and six in a pasture (Table 1; Fig. 1). The unequal number of sites per land-use category was due to limitations on accessibility. Each site consisted of two subsites, 1-3 km apart, to account for within-site variation. Each subsite consisted of six plots, placed at least 400 m apart and randomly distributed 80-100 m from a walking track.

**Table 1.**
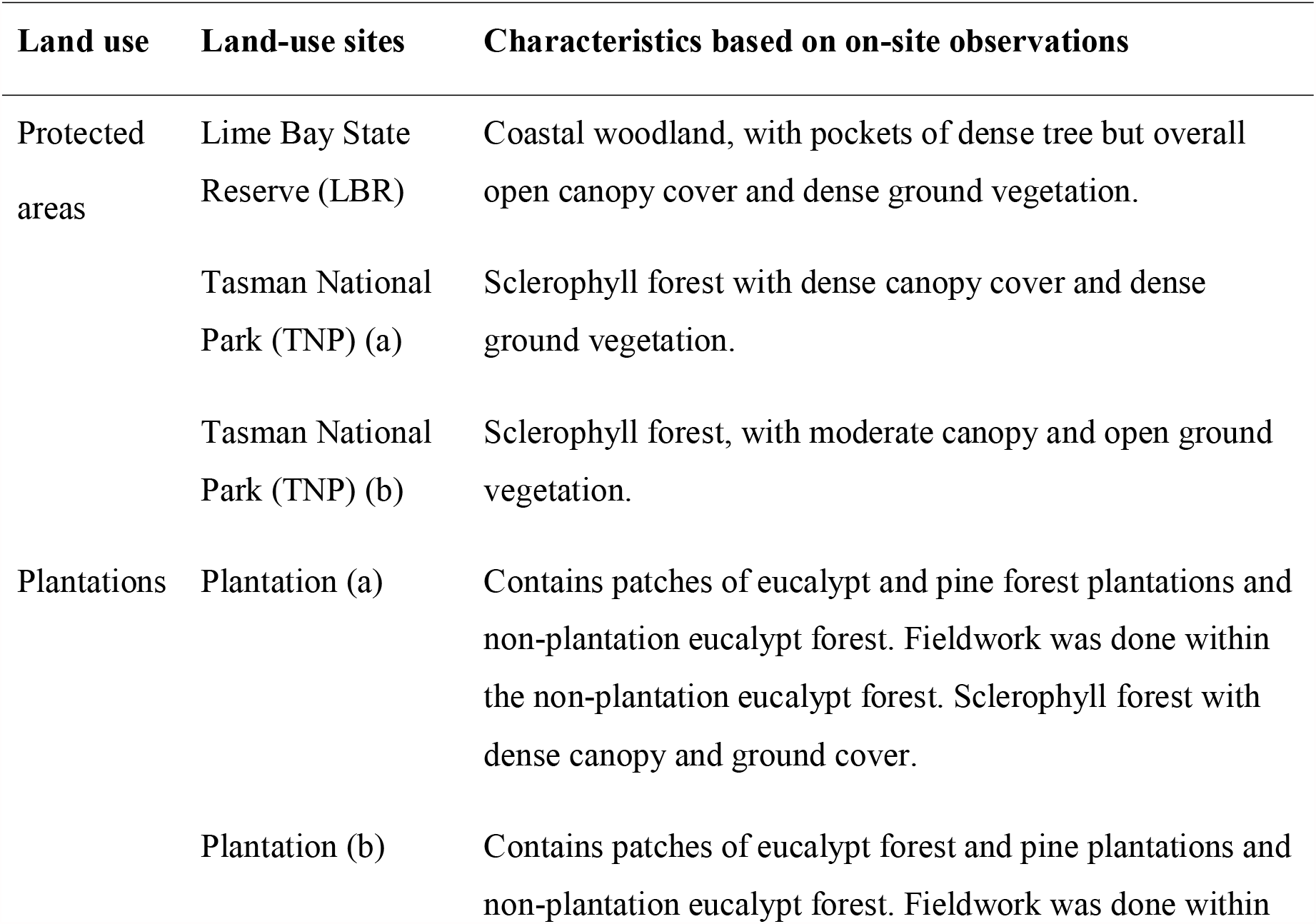
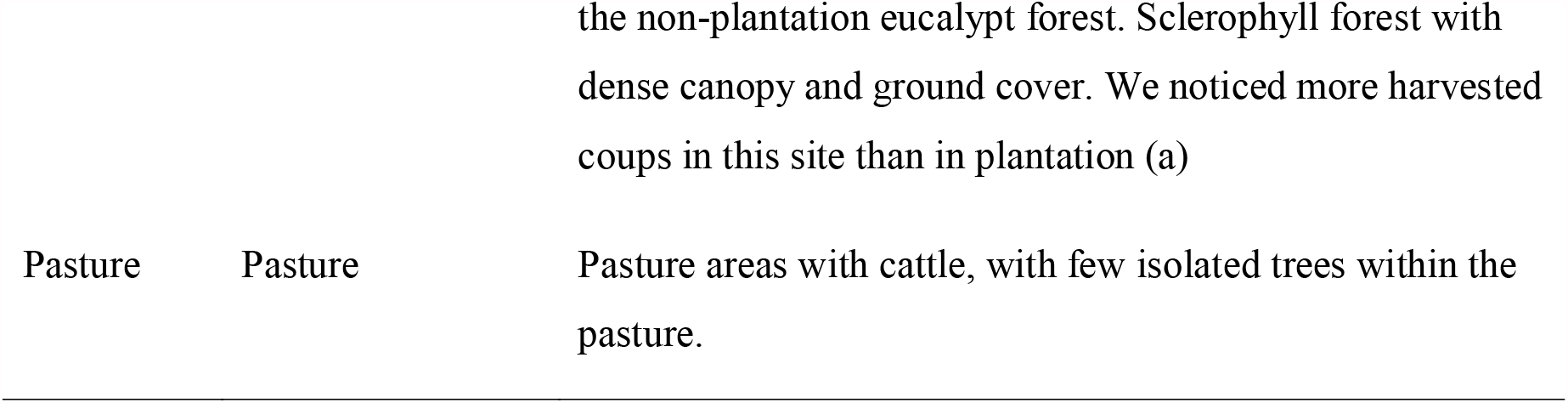
Characteristics of the different land-use sites within each land use

Within the 2-ha plots, we identified bird species and recorded their count visually and audibly following the standard protocol recommended by BirdLife (BirdLife 2021), which involves using a point-count method for 20 minutes. Honeyeaters (family *Meliphagidae*), were the only birds considered for this study, being the dominant group of nectivorous birds in Australia (Ford and Paton 1977). We also recorded the presence/absence of eucalyptus trees, which we identified to the species level using a field guide by Wiltshire and Potts (2007). The trees were grouped into the sub-genera *Symphyomyrtu*s and *Eucalyptus* (formerly *Monocalyptus*) (Nicolle 2015) as they have been shown to influence honeyeater presence (Dunkerley et al. 1990; Woinarski and Cullen 1984).

We established a total of 144 plots for bee and beetle observations by selecting, within each of the 36 × 2-ha plots, four different ground-level flowering plants. Where possible, four different species of flowering plant were chosen and if none was present no observation was made. Bees and beetles were sampled through visual observations, a commonly used technique to detect bee abundance (Prendergast et al. 2020). Bees were classified as either introduced or native; introduced bees being *Apis mellifera* (western honeybee) and *Bombus terrestris* (buff-tailed bumble bee), while native bees included all other bee species. Beetles were identified to family level using the iNaturalist website (https://inaturalist.org/). Flowering plants were identified to species or genus level using the University of Tasmania key to Tasmanian vascular plants (www.utas.edu.au/dicotkey/dicotkey/key.htm).

We established a total of 12 transects for butterfly observation. The transects were placed along the walking track between the three 2-ha plots in each of the land-use sites. There were six transects in protected areas (two in each of the three protected-areas land-use sites), four transects in plantations (two in each plantation) and two transects in pasture (Fig. 1). Butterflies were sampled along a 1000-m long and 5-m wide transect (Pollard 1977). An insect net was used to catch butterflies and record count and species. Species were photographed and identified using Common and Waterhouse (1972).

Observations took place between 07.00 – 11.00 h and 16.00 – 19.00 h for birds, and 10.00 – 15.00 h for insects, under low-wind and non-rainy conditions. Observations of all the investigated pollinator groups were repeated monthly during the Austral spring-summer of 2018, from September to December. The advantage of repeated sampling is that it yields higher precision within sites, at the cost of spatial bias, and importantly, it allowed us to account for any temporal changes in flowering vegetation and pollinator presence. Because of temporal changes in flowering vegetation and pollinator presence, we treated the repeated surveys as independent measurements. In total, we performed 144 surveys for birds (36 × 2-ha plots × 4 months), 576 surveys each for bees and beetles (36 × 2-ha plots × 4 flowering plants × 4 months) and 48 surveys for butterflies (12 transects × 4 months).

### Land-cover analysis

Land cover refers to the natural and artificial structures covering the land (Anderson et al. 1976); the land-cover classes considered in this study are ‘forest’ (areas dominated by trees, including plantations), ‘open’ (areas with low-lying vegetation, including shrubs and grasses), ‘barren’ (areas lacking vegetation), and ‘water’ (quantified as the percentage of total cover at different spatial scales). The allocation of land-cover classes was inferred from Sentinel 2 imagery as it has a high spatial resolution of 10 × 10 m (Sentinel Online 2021).

We carried out a pixel-based classification of the imagery following the method outlined by Diengdoh et al. (2020). This method involved, firstly, creating training and validation data for the above-mentioned land-cover classes in QGIS (QGIS Development Team 2021). The training and validation data were used for fitting and tuning the algorithms, and accuracy assessment, respectively. Secondly, machine-learning algorithms (hereafter algorithms) – support vector machine, Random Forests, k-nearest neighbour, and naïve Bayes were trained and used to classify the satellite imagery. Thirdly, the images were classified by the four algorithms using an unweighted ensemble algorithm (Diengdoh et al. 2020).

Accuracy was assessed by comparing the output to the validation data, which consisted of 100 points/pixels per land-cover class, randomly selected from the classified image and visually compared to imagery from Google Earth and field data for accuracy assessment. The output metrics we report include the overall accuracy (OA), the true-skill statistic (TSS) for the classified image, and the sensitivity and specificity of each land-cover class, where: OA is the number of correctly classified pixels divided by the total number of pixels examined (Foody 2002); TSS is equal to sensitivity plus specificity minus 1, where sensitivity is observed presences that are correctly predicted, and specificity is the observed absences that are predicted as such (Allouche et al. 2006). The classification analysis was done in R (R Core Team 2020).

The percentage of each land-cover class was calculated within buffers of two different sizes for each pollinator group. We used a buffer of a 2000 m radius (from the centre of the 2-ha plot and the 1000 m transect) to represent the land cover within the survey areas as well as its surrounding landscape, and a smaller buffer to represent the land cover within the survey area, which was 100 m in radius for the birds, bees, and beetles and 500 m for butterflies. We selected 500m for butterflies because a 100 m buffer did not encompass the entire 1000m transect. These buffer sizes are similar to those from other studies, e.g., birds (Smith et al. 2011), bees (Greenleaf et al. 2007) and butterflies (Bergman et al. 2004).

### Statistical analyses

We tested the ability of land use, land cover and plant genera to predict the abundance of each pollinator group. The multi-level categorical predictors ‘land use’ (3 levels) and ‘plant genera’ (24 levels), were transformed to dummy variables (i.e., binary presence/absence predictors) to reduce data dimensionality. Keeping land use and plant genera as constant predictors in the model, we compared a model including land cover within the survey area (100 or 500 m depending on taxa) with a model containing land cover within the surrounding area (2000 m).

Random Forests was used for predictive modelling, implemented using the *randomForest* R package (Liaw and Wiener 2002). The data was split into 70%/30% training/test using a random stratified method so that there was at least one repeated observation in the testing data that was not in the training data. The training data was used for model fitting, assessing partial dependence (PD) plots and individual conditional expectation (ICE) curves and calculating variable importance while the testing data was used for model accuracy, where the out-of-sample R^2^ and root mean square error (RMSE) of each model was calculated. Models were tuned for different *mtry* (the hyperparameter in Random Forests) values and the *mtry* value that resulted in the highest R^2^ was selected. The PD plots show the marginal effect of a predictor on the predicted outcome (Goldstein et al. 2015). The disadvantage of PD plots is that they show the average effect and heterogeneous effects may be hidden; we used ICE curves to assess those effects (Goldstein et al. 2015). The *pdp* r package (Greenwell 2017) was used for assessing PD plots and ICE curves.

## Results

### Pollinator richness and abundance

We observed a total of 297 honeyeaters belonging to eight different species, of which four are endemic to Tasmania (127 individuals); 511 bees, consisting of 284 native bees (184 *Exoneura* genus, 48 *Lasioglossum* genus and 52 individuals classified as others) and 227 introduced bees (211 honey bees and 16 bumble bees); 423 beetles belonging to nine families; and 84 butterflies belonging to eight species, of which only *Pieris rapae* is introduced. A list of species observed across the different land-use sites is included in Online Resource 1.

The median of the log count of different pollinator groups varied across the six LU sites (Fig. 2). Honeyeaters (birds) were more common across the three protected areas than in plantations or pastures (Fig. 2a). However, the median count of the insect pollinators was similar across the different LU sites (Fig. 2b-e).

**Fig. 2.**
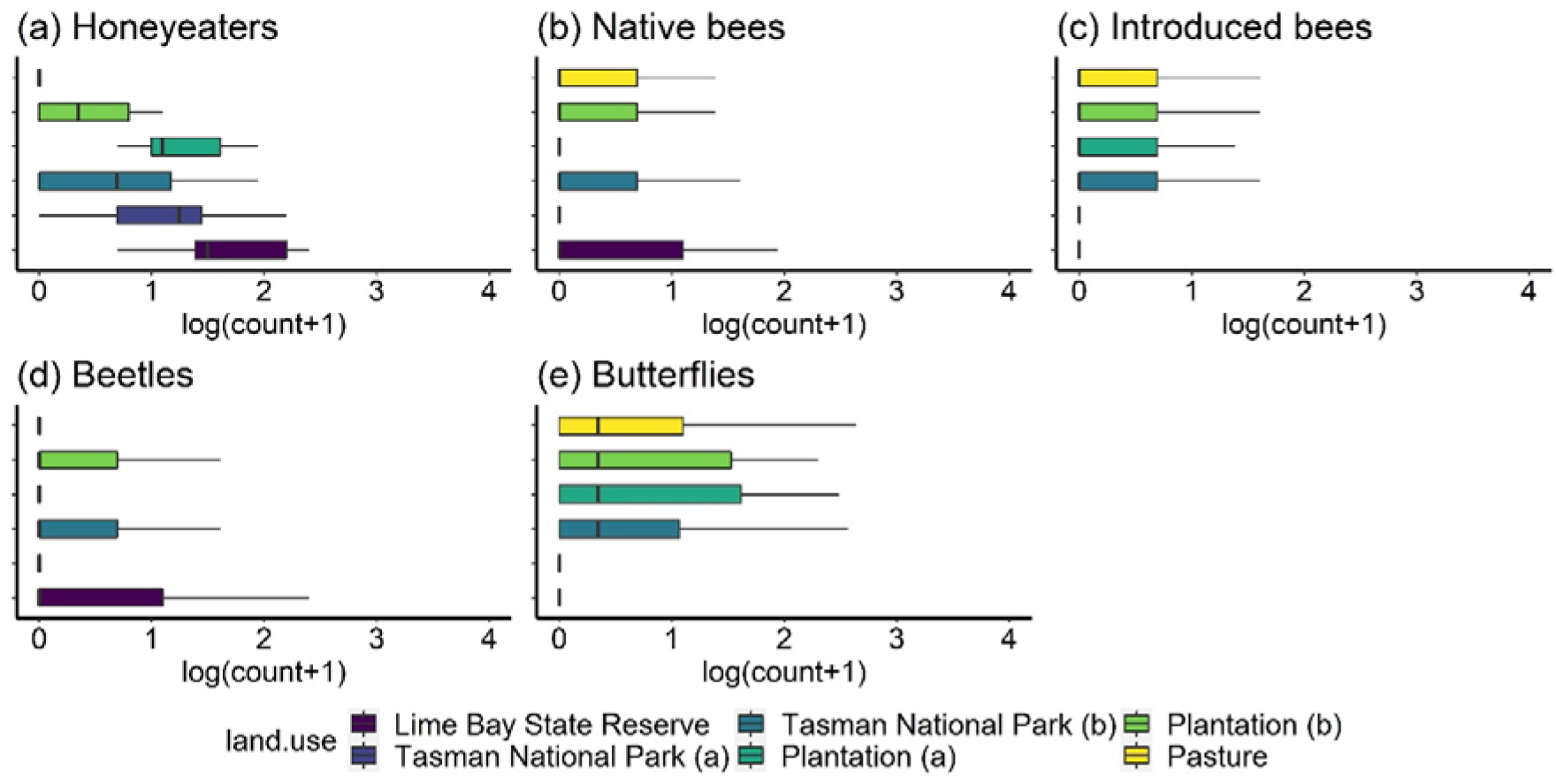
Log of total count (a) honeyeaters, (b-c) bees, (d) beetles and (e) butterflies across protected areas (Tasman National Park a and b, Lime Bay State Reserve), plantations (plantation a and b) and pastures land-use sites.

Pollinators showed preference for some plant genera. The median count of honeyeaters was higher in the presence of the eucalypt subgenus *Symphyomyrtus* compared with the subgenus *Eucalyptus* (Fig. 3a). Bees and beetles were observed visiting 24 plant genera, of which only three are exotic/naturalised (*Arctotheca, Taraxacum* and *Trifolium*, only found in pastures). The median count of native bees was highest in presence of *Pultenaea* and *Melaleuca* (Fig. 3b), while more introduced bees were found on *Anopterus, Lissanthe, Pimelea, Pomaderris*, and *Trifolium* flowers (Fig. 3c). The median count of beetles was highest in association with *Leptospermum, Pomaderris* and *Prostanthera* (Fig. 3d).

**Fig. 3.**
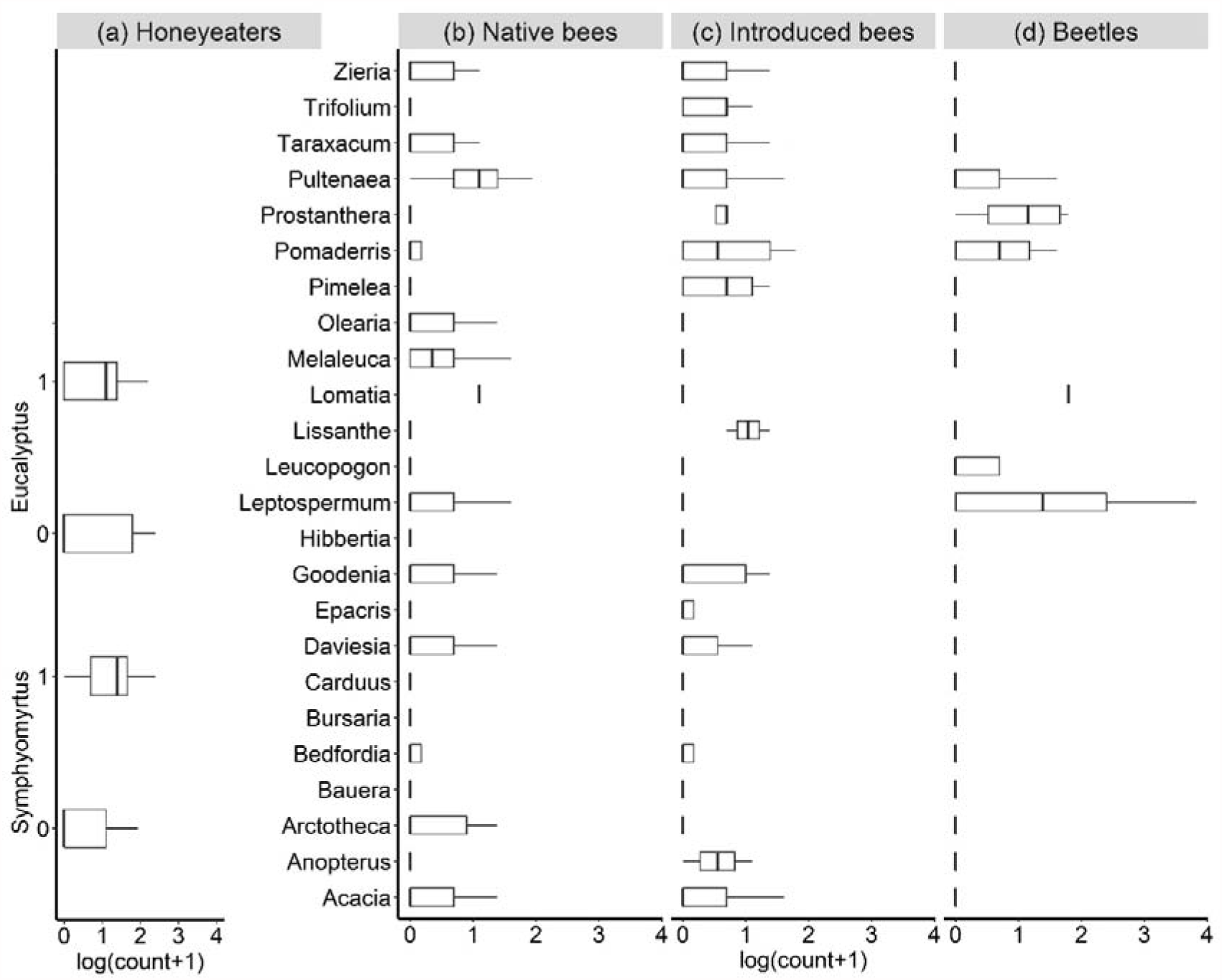
Log of total count (a) honeyeaters, (b) native bees, (c) introduced bees and (d) beetles per genus.

### Land-cover analysis

The classified image (Online Resource 2) had an overall accuracy of 79.5% with a 95% CI [75.2, 83.4] and a true skill statistic score of 77.4%. The confusion matrix of the classification results with sensitivity and specificity of the land-cover classes as well as the percentage of land cover within different buffers are included in the Online Resources (Online Resource 3-6).

### Predictors of pollinator count

The models including land cover within the survey area (100 or 500 m) performed better for introduced bees and beetles and explained 43.9% and 30.5% of the variation in pollinator abundance respectively, while the models containing land cover of the broader landscape (2000 m) were better for honeyeaters (49.1%) and native bees (21.6%; Table 2). The models for butterfly count were not usefully descriptive (0.01 R^2^; Online Resource 7). The land cover-forest within 2000 m buffer was the most important predictor for honeyeaters and native bees while open cover within 100 m buffer and *Leptospermum* were the most important predictor for introduced bees and beetles, respectively (Online Resource 8). The plant genera *Pultenaea* and *Acacia* were the most important predictors for native bees and introduced bees, respectively (Online Resource 8). Land use (i.e., human activities on the land) was not selected as the most important predictor for any of the models; however, it was consistently among the top 10 predictors for the different models (Online Resource 8).

**Table 2.**
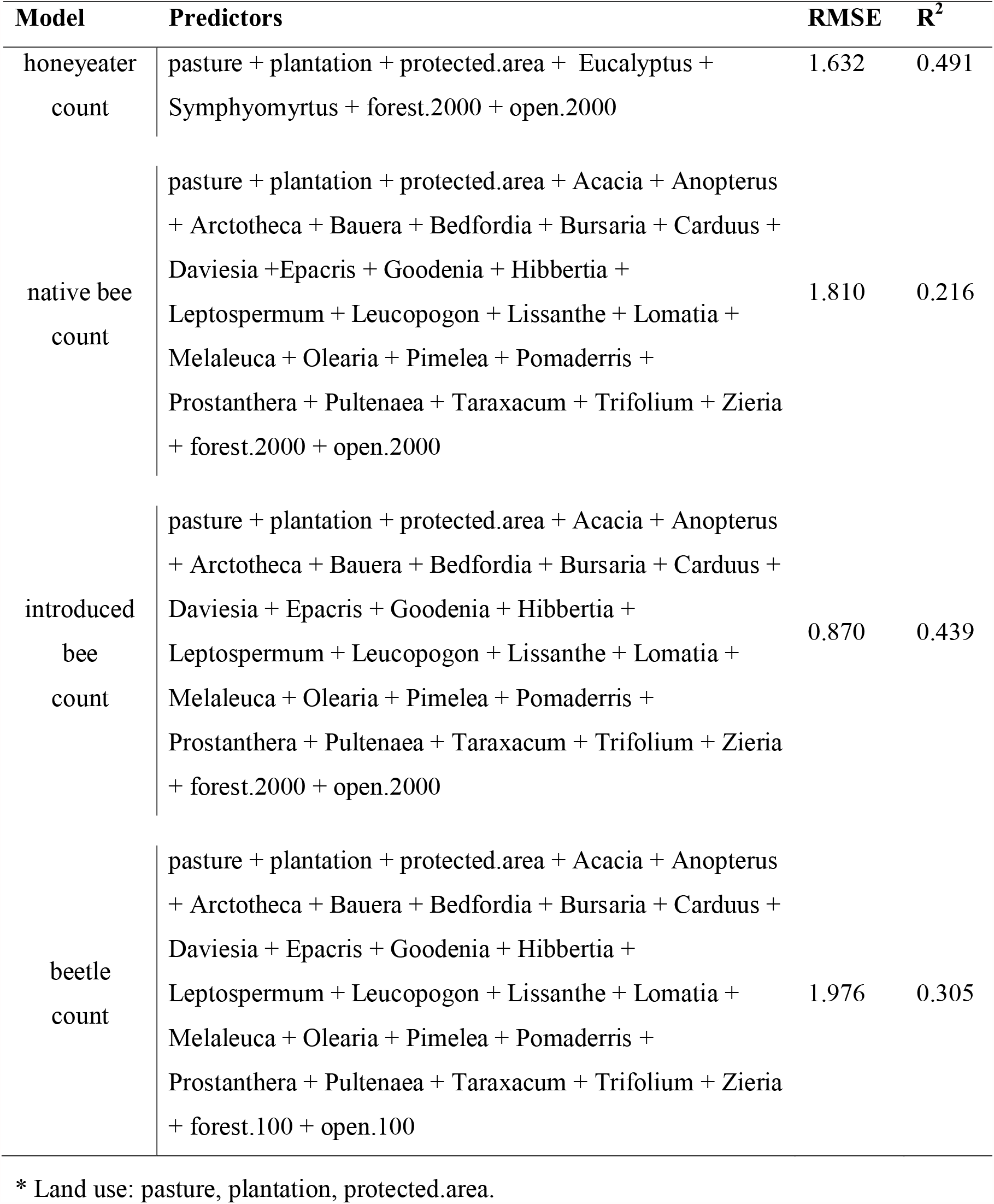
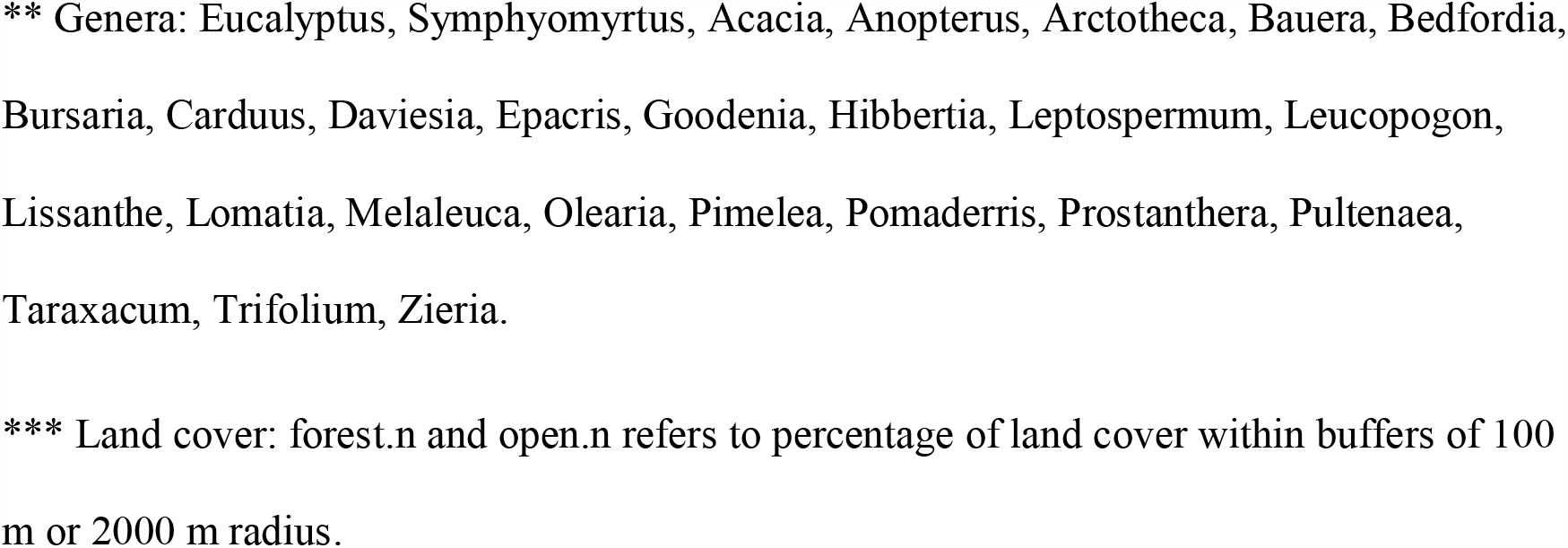
The predictive RMSE and R^2^ (i.e., on out-of-sample test data) of the different pollinator count models using the Random Forests algorithm.

| Model | Predictors | RMSE | $R^2$ |
| --- | --- | --- | --- |
| honeyeater<br>count | pasture + plantation + protected.area + Eucalyptus +<br>Symphyomyrtus + forest.2000 + open.2000 | 1.632 | 0.491 |
| native bee<br>count | pasture + plantation + protected.area + Acacia + Anopterus<br>+ Arctotheca + Bauera + Bedfordia + Bursaria + Carduus +<br>Daviesia + Epacris + Goodenia + Hibbertia +<br>Leptospermum + Leucopogon + Lissanthe + Lomatia +<br>Melaleuca + Olearia + Pimelea + Pomaderris +<br>Prostanthera + Pultenaea + Taraxacum + Trifolium + Zieria<br>+ forest.2000 + open.2000 | 1.810 | 0.216 |
| introduced<br>bee<br>count | pasture + plantation + protected.area + Acacia + Anopterus<br>+ Arctotheca + Bauera + Bedfordia + Bursaria + Carduus +<br>Daviesia + Epacris + Goodenia + Hibbertia +<br>Leptospermum + Leucopogon + Lissanthe + Lomatia +<br>Melaleuca + Olearia + Pimelea + Pomaderris +<br>Prostanthera + Pultenaea + Taraxacum + Trifolium + Zieria<br>+ forest.2000 + open.2000 | 0.870 | 0.439 |
| beetle<br>count | pasture + plantation + protected.area + Acacia + Anopterus<br>+ Arctotheca + Bauera + Bedfordia + Bursaria + Carduus +<br>Daviesia + Epacris + Goodenia + Hibbertia +<br>Leptospermum + Leucopogon + Lissanthe + Lomatia +<br>Melaleuca + Olearia + Pimelea + Pomaderris +<br>Prostanthera + Pultenaea + Taraxacum + Trifolium + Zieria<br>+ forest.100 + open.100 | 1.976 | 0.305 |

\* Land use: pasture, plantation, protected.area.
\*\* Genera: Eucalyptus, Symphyomyrtus, Acacia, Anopterus, Arctotheca, Bauera, Bedfordia, Bursaria, Carduus, Daviesia, Epacris, Goodenia, Hibbertia, Leptospermum, Leucopogon, Lissanthe, Lomatia, Melaleuca, Olearia, Pimelea, Pomaderris, Prostanthera, Pultenaea, Taraxacum, Trifolium, Zieria.
\*\*\* Land cover: forest.n and open.n refers to percentage of land cover within buffers of 100 m or 2000 m radius.

The effect land use had on pollinator count varied between and within pollinator taxa. Protected areas had a positive effect on honeyeater count, but on average had no effect on bees or beetle count (Fig. 4a, d, g, j). However, based on the ICE plots, we found protected areas had both positive and negative effects on bees and beetles (Fig. 4d, g, j). Plantations had a positive effect on beetles but no effect on honeyeater and bees count (Fig. 4b, e, h, k). Yet, based on the ICE plots, we found plantations had both positive and negative effect on honeyeater and bees count (Fig. 4b, e, h). Pasture was the only land use which had a negative effect on honeyeater and beetle count (Fig. 4c, l).

**Fig. 4.**
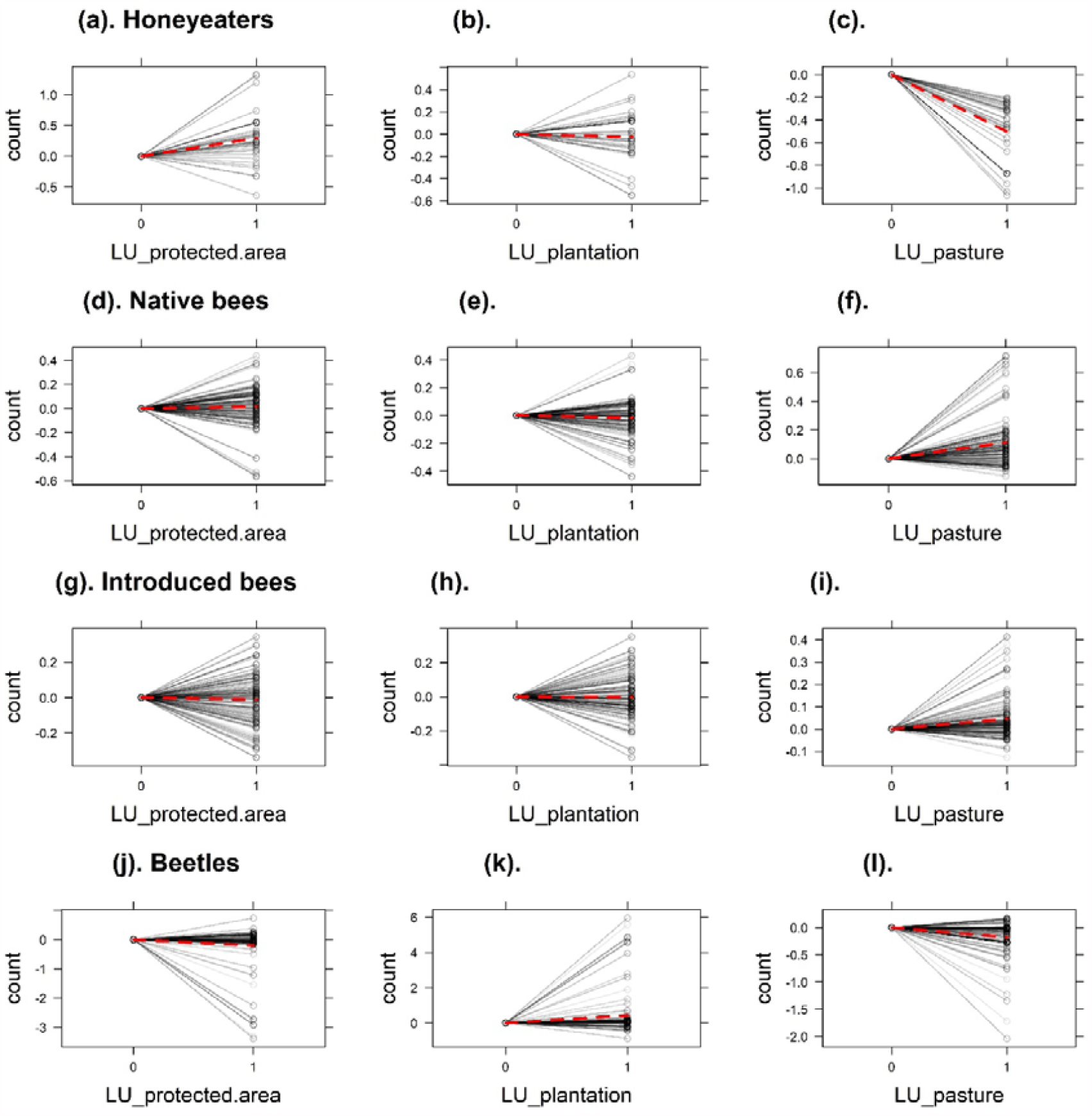
Partial dependence plots (dashed-red lines) and individual conditional expectation (ICE; solid-black lines) curves of the land-use predictors on (a-c) honeyeaters, (d-f) native bees, (g-i) introduced bees, and (j-l) beetle models.

Land cover did not have a monotonic or linear relationship with the number of pollinators; rather, it exhibited a stepwise relation with pollinator abundance (Fig. 5). The percentage of forest cover had a positive effect on the number of honeyeaters (Fig. 5b). Conversely, open cover had a negative effect on honeyeaters but positive effect on bees (Fig. 5a, c, e). Overall, the presence of the different plant genera had a positive effect on the number of different pollinator groups, except for *Eucalyptus* subgenus *Eucalyptus*, which had a negative effect on honeyeater count (Fig. 6a). *Leucopogon, Pultenaea* and *Acacia* were the three most important variables for native-bee abundance, while *Acacia, Melaleuca*, and *Pomaderris* were the three most important variables for introduced bees (Online Resource 8; Fig. 6c-h). *Leptospermum Pultenaea* and *Pomaderris* were most important for beetle abundance (Online Resource 8; Fig. 6. i-k).

**Fig. 5.**
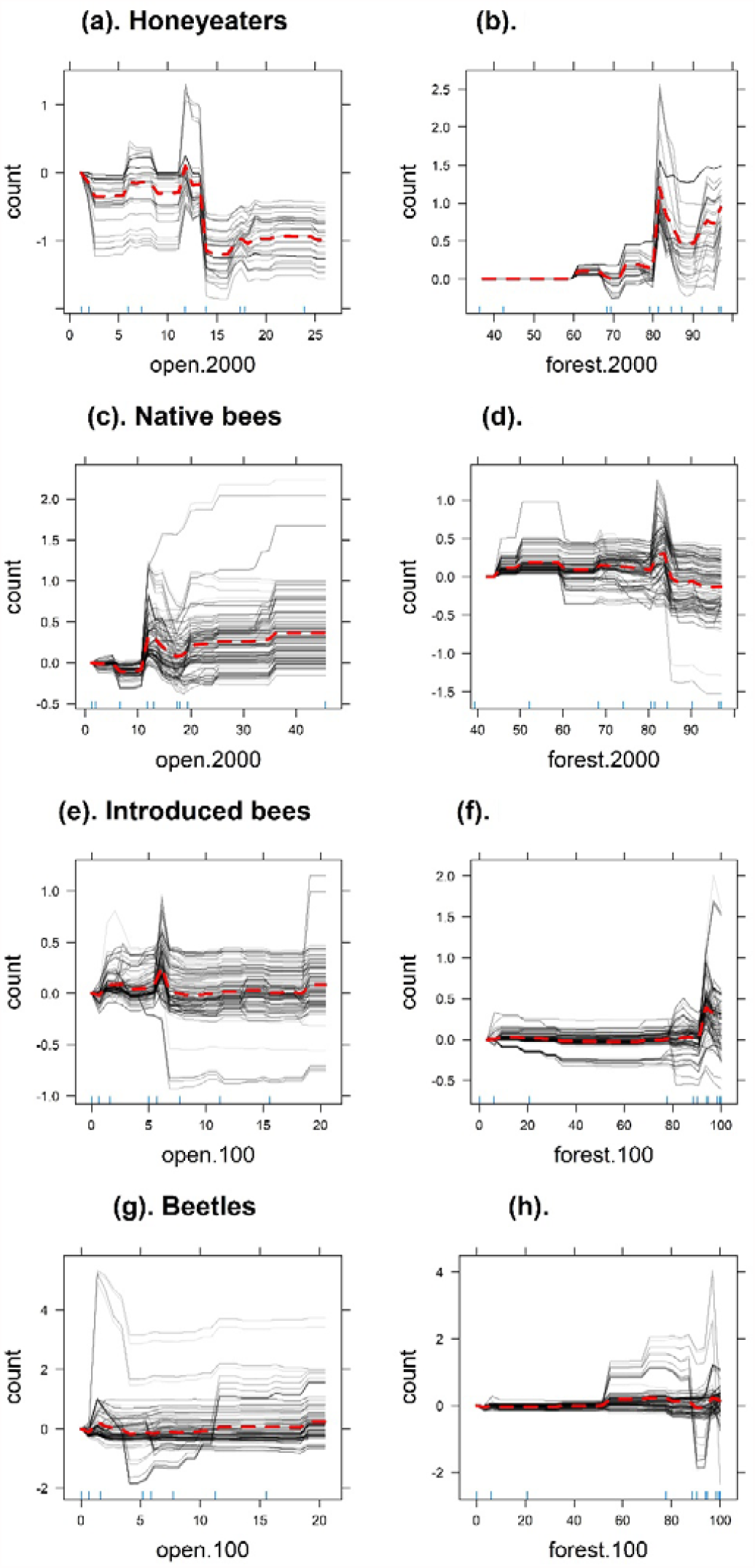
Partial dependence plots (dashed-red lines) and individual conditional expectation (ICE; solid-black lines) curves of the land-cover predictors for (a-b) honeyeaters, (c-d) native bees, (e-f) introduced bees, and (g-h) beetle models.

**Fig. 6.**
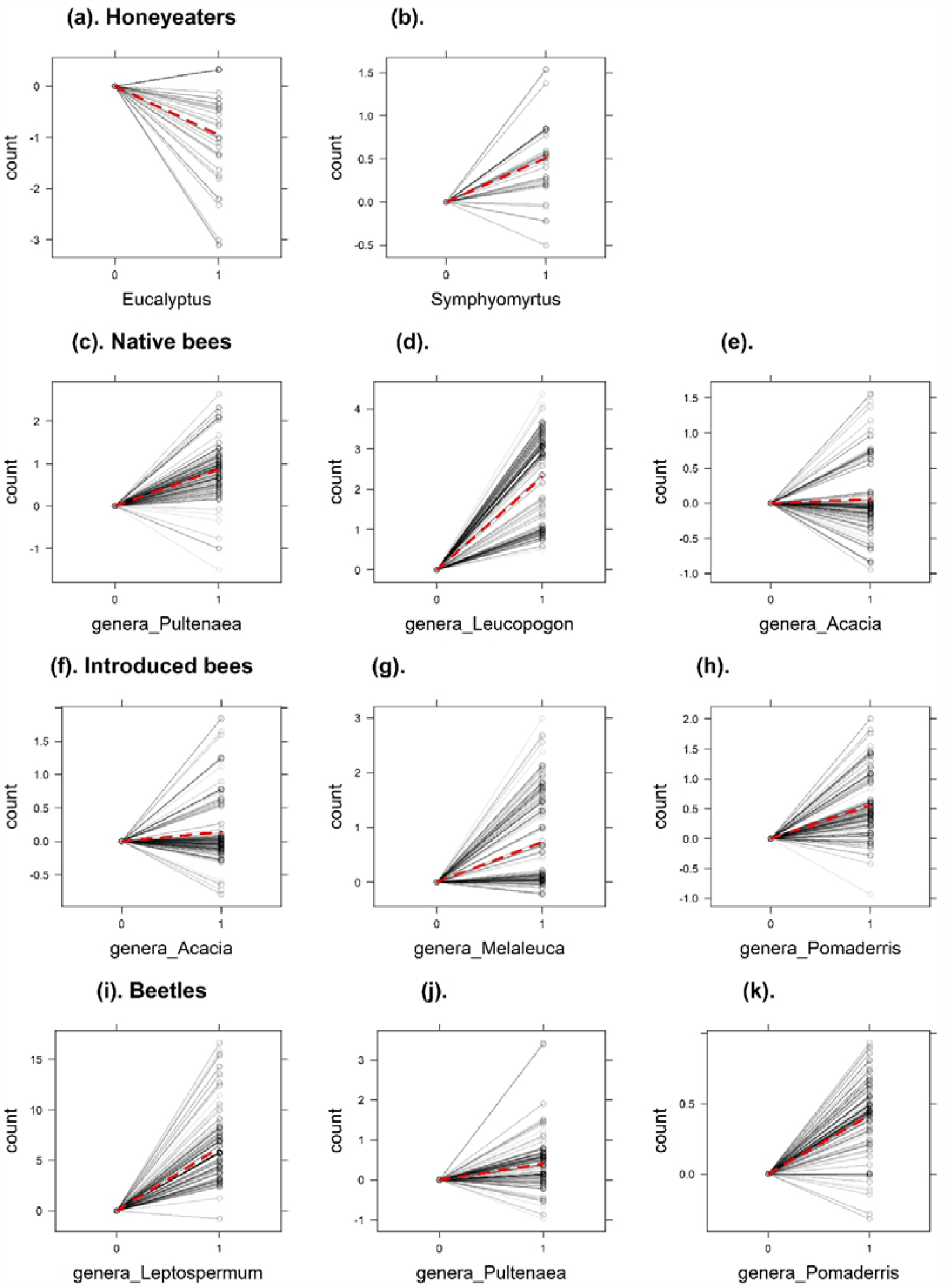
Partial dependence plots (dashed-red lines) and individual conditional expectation (ICE; solid-black lines) curves of the most important genera predictors for (a-b) honeyeaters, (c-e) native bees, (f-h) introduced bees, and (i-k) beetle models.

## Discussion

We assessed the effect of land use, land cover, and plant genera, on the abundance of multiple pollinator groups across a mixed-use, moderately human-modified landscape. We found the percentage of land cover type i.e., forest versus open land cover was the most important predictor for all models as land cover serves as a source of foraging and nesting (Kremen et al. 2007; Öckinger et al. 2012). As expected, given the dependence pollinators on pollen and nectar, the presence of some plant genera had a positive effect on pollinator abundance. The effect of land use on pollinator abundance varied depending on the pollinating taxa, consistent with previous findings (Winfree et al. 2011).

The lower importance of land use compared to land cover and plant genera is consistent with existing literature (e.g., De Palma et al. 2016) and suggests that land use might become important only when extremely modified land-use types (e.g., pure agriculture, or urban) are considered (Millard et al. 2021; Winfree et al. 2009; Winfree et al. 2011). The availability of natural habitat and floral resources present in and around different land-use types are more likely to affect pollinators; for example, (1) pollinators can make use of the pollen and nectar provided by plants in road verges or green infrastructure in agricultural and urban landscapes (Daniels et al. 2020; Phillips et al. 2019) and (2) agricultural landscapes with higher amount of high-quality habitat support a higher abundance and richness of pollinators than agricultural landscapes without such habitat (Kavanagh et al. 2007; Kennedy et al. 2013). The positive effect of plantation and pasture land-use types on pollinator taxa could be attributed to the low land-use intensity of the overall study area (Millard et al. 2021) and/or, due to the availability of plantation and non-plantation forest cover and floral resources in plantations and pastures, respectively (Brito et al. 2017; Kavanagh et al. 2007; Kennedy et al. 2013).

The between-taxa variation on the effect of land use was likely due to species-specific requirements and traits, and the within-taxa variation could be due to the coarse taxonomic identification we used. Species traits are known to influence pollinator response to anthropogenic factors (Cariveau and Winfree 2015; De Palma et al. 2015). The visual sampling of bees and beetles allowed us to assess which plants they visited but limited the taxonomic identification of insects, and small samples sizes made the testing of species-specific responses to the predictor variables unfeasible.

The predictive capacity of land cover within the survey area (100 m buffer) versus the land cover within the surrounding area (2000 m buffer) depended on the pollinating taxon, which is consistent with the literature (Bartholomée et al. 2020; Kennedy et al. 2013; Steffan-Dewenter et al. 2002). A possible explanation is that birds, being more mobile, can cover larger distances, and as a result the surrounding land cover was a better predictor than the survey land cover. The difference been introduced bees and native bees could be due to their sizes. Introduced bees consist of honey bees and bumble bees, which are comparatively larger than the native bees. For pollinators such as bees, body size is considered an important factor in their response to land cover at both local and landscape scales (Benjamin et al. 2014). The spatial and temporal distribution of resources is another factor that can influence the effect of land-cover scale on pollinators (Pufal et al. 2017). For instance, the negative impact of open cover on honeyeaters could be due to the lack of resources, as open areas—which tend to lack resources such as food and shelter—are known to negatively impact honeyeaters (Kavanagh et al. 2007), whereas forest cover has been shown to positively influence them (Harrisson et al. 2014). By contrast, resources required by both native and introduced bees could be found in open areas, hence the positive effect on those taxa, consistent with the findings from previous studies (Kaluza et al. 2016). The ICE plots of both open and forest cover showed both had positive and negative impacts on beetles, which might be due to micro-climate, leaf litter and soil variables, all being known to influence beetles (Fountain-Jones et al. 2015). This suggests that the overall low effect of land cover on beetle counts could be due simply to data paucity. The stepwise relation between land cover and pollinator count could be due to a lack of data covering the entire gradient of percentage of land cover, which would result in the effect ‘peaking’ or ‘dipping’ at only certain percentages.

The presence of the eucalypt subgenus *Eucalyptus* was associated with lower honeyeater counts, possibly because of the high abundance of arthropods associated with subgenus *Symphyomyrtus* compared with *Eucalyptus* (Dunkerley et al. 1990; Saunders and Burgin 2001). This could have led honeyeaters, particularly insectivorous species such as Yellow-throated, Strong-billed, and Black-headed honeyeaters (Thomas 1980), to be more attracted to *Symphyomyrtus* trees. Pollinator insect abundance was also impacted by the presence of the different plant genera. In particular, the presence of the native genera *Acacia, Pultenaea, Pomaderris, Leucopogon, Melaleuca*, and *Leptospermum* was associated with more native bees, introduced bees, and beetles. All these relationships have plausible biological underpinnings. For example, *Acacia* produces copious pollen (Stone et al. 2003) which is essential for larval provisions for almost all bees (Brosi et al. 2007) while *Pultenaea* and *Pomaderris* contain nectar and pollen and are pollinated by bees, beetles and butterflies (Armstrong 1979; De Kok and West 2004; Ogilvie et al. 2009). *Leptospermum*, in addition to providing nectar and pollen, is also a rich source of fruits and grass-root material which beetles and their larvae feed upon (Andersen and New 1987; Stephens et al. 2005).

Overall, we did fond that one size does not fit all—that is, the effects and predictive capacity of land use and land cover in a mixed-use landscape with moderate changes varied depending on the pollinating taxa. Indeed, our results highlight the complexity of pollinator-landscape interactions and that different taxa require different conservation and management policies.

Based on our results for the Tasman Peninsula, we recommend the conservation of a mosaic of forest and open land cover, at both small and large scales. We also recommend the management of native plants (and particularly species belonging to the *Acacia, Leucopogon, Leptospermum, Melaleuca, Pomaderris*, and *Pultenaea* genera and the subgenus *Symphyomyrtus* in Tasmania and mainland Australia where applicable) to attract and sustain a community of pollinators.

More generally (beyond this case study), such taxon-specific requirements would be of particular importance in landscapes with extreme change or those dominated by a single land-use type (e.g., urban areas), where creating a mosaic of land-cover types is not an achievable option. These recommendations fall within a known framework of choosing native plants that provide nectar and/or pollen resources for a long temporal span (M’Gonigle et al. 2017; Menz et al. 2011). A major advantage of the approach taken in this study, involving machine learning and predictive modelling, is in its ability to subject conservation-management problems to robust, data-driven assessments, and use these to make informed decisions (Thessen 2016; Willcock et al. 2018). Future studies should consider using both active and passive sampling techniques as well as human observations and mechanised methods such as acoustic recorders for birds (Wimmer et al. 2013) and photographic recording of bees (Steen 2017), to obtain a more detailed representation of pollinator communities. Although a past-present comparison would be particularly beneficial for assessing the impacts of land-use and land-cover changes on pollinators, we lacked baseline data on pollinators, making comparative space-for-time assessments a logical substitute. Future studies should explore landscapes and databases that provide opportunities to make past-present comparisons, to better understand the impact of land-use and land-cover changes on pollinator taxa.

## Supporting information

Online Resources

## Acknowledgements

This work was supported by the Australian Research Council [grant number FL160100101]. The authors acknowledge the Pydairrerme people, traditional custodians of the land where fieldwork was carried out. We thank the different landowners for the permission provided to carry out fieldwork on their properties. We would also like to thank the volunteers who assisted with fieldwork.

## References

Allouche O, Tsoar A, Kadmon R (2006) Assessing the accuracy of species distribution models: prevalence, kappa and the true skill statistic (TSS). Journal of Applied Ecology 43:1223–1232. https://doi.org/10.1111/j.1365-2664.2006.01214.x

Andersen AN, New T (1987) Insect inhabitants of fruits of Leptospermum, Eucalyptus and Casuarina in southeastern Australia. Australian Journal of Zoology 35:327–336. https://doi.org/10.1071/ZO9870327

Anderson JR, Hardy EE, Roach JT, Witmer RE (1976) A land use and land cover classification system for use with remote sensor data. Professional Paper 964. US Government Printing Office, Washington DC https://doi.org/10.3133/pp964

Armstrong J (1979) Biotic pollination mechanisms in the Australian flora—a review. New Zealand Journal of Botany 17:467–508. https://doi.org/10.1080/0028825X.1979.10432565

Australian Bureau of Statistics (2021) Region summary: Tasman (M). https://dbr.abs.gov.au/region.html?lyr=lga&rgn=65210. Accessed 22/04/2021

Bartholomée O, Aullo A, Becquet J, Vannier C, Lavorel S (2020) Pollinator presence in orchards depends on landscape-scale habitats more than in-field flower resources. Agriculture, Ecosystems & Environment 293:106806. https://doi.org/10.1016/j.agee.2019.106806

Baude M, Kunin WE, Boatman ND, Conyers S, Davies N, Gillespie MA, Morton RD, Smart SM, Memmott J (2016) Historical nectar assessment reveals the fall and rise of floral resources in Britain. Nature 530:85. https://doi.org/10.1038/nature16532

Benjamin FE, Reilly JR, Winfree R (2014) Pollinator body size mediates the scale at which land use drives crop pollination services. Journal of Applied Ecology 51:440–449. https://doi.org/10.1111/1365-2664.12198

Bergman KO, Askling J, Ekberg O, Ignell H, Wahlman H, Milberg P (2004) Landscape effects on butterfly assemblages in an agricultural region. Ecography 27:619–628. https://doi.org/10.1111/j.0906-7590.2004.03906.x

BirdLife (2021) Survey Techniques. https://birdata.birdlife.org.au/survey-techniques. Accessed 22/04/2021

Brito TF, Phifer CC, Knowlton JL, Fiser CM, Becker NM, Barros FC, Contrera FA, Maués MM, Juen L, Montag LF (2017) Forest reserves and riparian corridors help maintain orchid bee (Hymenoptera: Euglossini) communities in oil palm plantations in Brazil. Apidologie 48:575–587. https://doi.org/10.1007/s13592-017-0500-z

Brosi BJ, Daily GC, Ehrlich PR (2007) Bee community shifts with landscape context in a tropical countryside. Ecological Applications 17:418–430. https://doi.org/10.1890/06-0029

Brown M, Duncan F (1986) The vegetation of Tasman Peninsula. Papers and Proceedings of the Royal Society of Tasmania:33–50.

Campbell AJ, Carvalheiro LG, Gastauer M, Almeida-Neto M, Giannini TC (2019) Pollinator restoration in Brazilian ecosystems relies on a small but phylogenetically-diverse set of plant families. Scientific Reports 9:1–10. https://doi.org/10.1038/s41598-019-53829-4

Cariveau DP, Winfree R (2015) Causes of variation in wild bee responses to anthropogenic drivers. Current Opinion in Insect Science 10:104–109. https://doi.org/10.1016/j.cois.2015.05.004

Common IFB, Waterhouse DF (1972) Butterflies of Australia. Angus and Robertson, Sydney

Daniels B, Jedamski J, Ottermanns R, Ross-Nickoll M (2020) A “plan bee” for cities: Pollinator diversity and plant-pollinator interactions in urban green spaces. PLoS One 15:e0235492. https://doi.org/10.1371/journal.pone.0235492

De Kok R, West J (2004) A revision of the genus Pultenaea (Fabaceae). 3. The eastern species with recurved leaves. Australian Systematic Botany 17:273–326. https://doi.org/10.1071/SB02028

De Palma A, Abrahamczyk S, Aizen MA, Albrecht M, Basset Y, Bates A, Blake RJ, Boutin C, Bugter R, Connop S (2016) Predicting bee community responses to land-use changes: Effects of geographic and taxonomic biases. Scientific Reports 6:31153. https://doi.org/10.1038/srep31153

De Palma A, Kuhlmann M, Roberts SP, Potts SG, Börger L, Hudson LN, Lysenko I, Newbold T, Purvis A (2015) Ecological traits affect the sensitivity of bees to land□use pressures in European agricultural landscapes. Journal of Applied Ecology 52:1567–1577. https://doi.org/10.1111/1365-2664.12524

Department of Primary Industries Parks Water and Environment (2015) Tasmanian Land Use 2015.

Diengdoh VL, Ondei S, Hunt M, Brook BW (2020) A validated ensemble method for multinomial land-cover classification. Ecological Informatics 56:101065. https://doi.org/10.1016/j.ecoinf.2020.101065

Dunkerley GM, Ford H, Danthanarayana W (1990) The fuscous honeyeater: food resources and the bird community. Dissertation, University of New England

Foody GM (2002) Status of land cover classification accuracy assessment. Remote Sensing of Environment 80:185–201. https://doi.org/10.1016/S0034-4257(01)00295-4

Ford HA, Paton DC (1977) The comparative ecology of ten species of honeyeaters in South Australia. Australian Journal of Ecology 2:399–407. https://doi.org/10.1111/j.1442-9993.1977.tb01155.x

Fountain-Jones NM, Jordan GJ, Baker TP, Balmer JM, Wardlaw T, Baker SC (2015) Living near the edge: being close to mature forest increases the rate of succession in beetle communities. Ecological Applications 25:800–811. https://doi.org/10.1890/14-0334.1

Goldstein A, Kapelner A, Bleich J, Pitkin E (2015) Peeking inside the black box: Visualizing statistical learning with plots of individual conditional expectation. Journal of Computational and Graphical Statistics 24:44–65. https://doi.org/10.1080/10618600.2014.907095

Greenleaf SS, Williams NM, Winfree R, Kremen C (2007) Bee foraging ranges and their relationship to body size. Oecologia 153:589–596. https://doi.org/10.1007/s00442-007-0752-9

Greenwell BM (2017) pdp: An R Package for Constructing Partial Dependence Plots. The R Journal 9:421–436. https://journal.r-project.org/archive/2017/RJ-2017-016/index.html

Harrisson KA, Pavlova A, Amos JN, Radford JQ, Sunnucks P (2014) Does reduced mobility through fragmented landscapes explain patch extinction patterns for three honeyeaters? Journal of Animal Ecology 83:616–627. https://doi.org/10.1111/1365-2656.12172

Kaluza BF, Wallace H, Heard TA, Klein AM, Leonhardt SD (2016) Urban gardens promote bee foraging over natural habitats and plantations. Ecology and Evolution 6:1304–1316. https://doi.org/10.1002/ece3.1941

Kavanagh RP, Stanton MA, Herring MW (2007) Eucalypt plantings on farms benefit woodland birds in south□eastern Australia. Austral Ecology 32:635–650.

Kennedy CM, Lonsdorf E, Neel MC, Williams NM, Ricketts TH, Winfree R, Bommarco R, Brittain C, Burley AL, Cariveau D (2013) A global quantitative synthesis of local and landscape effects on wild bee pollinators in agroecosystems. Ecology Letters 16:584–599. https://doi.org/10.1111/ele.12082

Klein AM, Vaissiere BE, Cane JH, Steffan-Dewenter I, Cunningham SA, Kremen C, Tscharntke T (2007) Importance of pollinators in changing landscapes for world crops. Proceedings of the Royal Society B: Biological Sciences 274:303–313. https://doi.org/10.1098/rspb.2006.3721

Kral-O’Brien KC, O’Brien PL, Hovick TJ, Harmon JP (2021) Meta-analysis: Higher Plant Richness Supports Higher Pollinator Richness Across Many Land Use Types. Annals of the Entomological Society of America https://doi.org/10.1093/aesa/saaa061

Kremen C, Williams NM, Aizen MA, Gemmill□Herren B, LeBuhn G, Minckley R, Packer L, Potts SG, Roulston Ta, Steffan□Dewenter I (2007) Pollination and other ecosystem services produced by mobile organisms: a conceptual framework for the effects of land□use change. Ecology Letters 10:299–314. https://doi.org/10.1111/j.1461-0248.2007.01018.x

Liaw A, Wiener M (2002) Classification and Regression by randomForest. R News 2:18–22. https://CRAN.R-project.org/doc/Rnews/

M’Gonigle LK, Williams NM, Lonsdorf E, Kremen C (2017) A Tool for Selecting Plants When Restoring Habitat for Pollinators. Conservation Letters 10:105–111. https://doi.org/10.1111/conl.12261

Maes D, Van Dyck H (2001) Butterfly diversity loss in Flanders (north Belgium): Europe’s worst case scenario? Biological Conservation 99:263–276. https://doi.org/10.1016/S0006-3207(00)00182-8

Menz MH, Phillips RD, Winfree R, Kremen C, Aizen MA, Johnson SD, Dixon KW (2011) Reconnecting plants and pollinators: challenges in the restoration of pollination mutualisms. Trends in Plant Science 16:4–12. https://doi.org/10.1016/j.tplants.2010.09.006

Millard J, Outhwaite CL, Kinnersley R, Freeman R, Gregory RD, Adedoja O, Gavini S, Kioko E, Kuhlmann M, Ollerton J (2021) Global effects of land-use intensity on local pollinator biodiversity. Nature Communications 12:1–11. https://doi.org/10.1038/s41467-021-23228-3

Montero-Castaño A, Vilà M (2012) Impact of landscape alteration and invasions on pollinators: a meta-analysis. Journal of Ecology 100:884–893. https://doi.org/10.1111/j.1365-2745.2012.01968.x

Nicolle D (2015) Classification of the eucalypts (Angophora, Corymbia and Eucalyptus) Version 4. https://dn.com.au/Classification-Of-The-Eucalypts.pdf.

Nicolson SW, Wright GA (2017) Plant–pollinator interactions and threats to pollination: perspectives from the flower to the landscape. Functional Ecology 31:22–25. https://doi.org/10.1111/1365-2435.12810

Öckinger E, Bergman K-O, Franzén M, Kadlec T, Krauss J, Kuussaari M, Pöyry J, Smith HG, Steffan-Dewenter I, Bommarco R (2012) The landscape matrix modifies the effect of habitat fragmentation in grassland butterflies. Landscape Ecology 27:121–131. https://doi.org/10.1007/s10980-011-9686-z

Ogilvie JE, Zalucki JM, Boulter SL (2009) Pollination biology of the sclerophyllous shrub Pultenaea villosa Willd.(Fabaceae) in southeast Queensland, Australia. Plant Species Biology 24:11–19. https://doi.org/10.1111/j.1442-1984.2009.00235.x

Ollerton J, Erenler H, Edwards M, Crockett R (2014) Pollinator declines. Extinctions of aculeate pollinators in Britain and the role of large-scale agricultural changes. Science 346:1360–1362. https://doi.org/10.1126/science.1257259

Ollerton J, Winfree R, Tarrant S (2011) How many flowering plants are pollinated by animals? Oikos 120:321–326. https://doi.org/10.1111/j.1600-0706.2010.18644.x

Paton DC (2000) Disruption of bird□plant pollination systems in southern Australia. Conservation Biology 14:1232–1234. https://doi.org/10.1046/j.1523-1739.2000.00015.x

Phillips BB, Gaston KJ, Bullock JM, Osborne JL (2019) Road verges support pollinators in agricultural landscapes, but are diminished by heavy traffic and summer cutting. Journal of Applied Ecology 56:2316–2327. https://doi.org/10.1111/1365-2664.13470

Pollard E (1977) A method for assessing changes in the abundance of butterflies. Biological Conservation 12:115–134. https://doi.org/10.1016/0006-3207(77)90065-9

Potts SG, Biesmeijer JC, Kremen C, Neumann P, Schweiger O, Kunin WE (2010) Global pollinator declines: trends, impacts and drivers. Trends in Ecology & Evolution 25:345–353. https://doi.org/10.1016/j.tree.2010.01.007

Prendergast KS, Menz MHM, Dixon KW, Bateman PW (2020) The relative performance of sampling methods for native bees: an empirical test and review of the literature. Ecosphere 11:e03076. https://doi.org/10.1002/ecs2.3076

Pufal G, Steffan-Dewenter I, Klein A-M (2017) Crop pollination services at the landscape scale. Current Opinion in Insect Science 21:91–97. https://doi.org/10.1016/j.cois.2017.05.021

QGIS Development Team (2021) QGIS Geographic Information System. QGIS Association. version 3.12.1. https://www.qgis.org

R Core Team (2020) R: A language and environment for statistical computing. R Foundation for Statistical Computing, Vienna, Austria. version 4.0.5. https://www.R-project.org/

Regan EC, Santini L, Ingwall-King L, Hoffmann M, Rondinini C, Symes A, Taylor J, Butchart SHM (2015) Global Trends in the Status of Bird and Mammal Pollinators. Conservation Letters 8:397–403. https://doi.org/10.1111/conl.12162

Sabatino M, Rovere A, Meli P (2021) Restoring pollination is not only about pollinators: Combining ecological and practical information to identify priority plant species for restoration of the Pampa grasslands of Argentina. Journal for Nature Conservation 61:126002. https://doi.org/10.1016/j.jnc.2021.126002

Saunders AS, Burgin S (2001) Selective foliage foraging by Red Wattlebirds, Anthochaera carunculata, and Noisy Friarbirds, Philemon corniculatus. Emu 101:163–166. https://doi.org/10.1071/MU00007

Senapathi D, Goddard MA, Kunin WE, Baldock KCR (2016) Landscape impacts on pollinator communities in temperate systems: evidence and knowledge gaps. Functional Ecology 31:26–37. https://doi.org/10.1111/1365-2435.12809

Sentinel Online (2021) Sentinel-2 Spatial Resolutions https://sentinel.esa.int/web/sentinel/user-guides/sentinel-2-msi/resolutions/spatial. Accessed 22/04/2021

Smith AC, Fahrig L, Francis CM (2011) Landscape size affects the relative importance of habitat amount, habitat fragmentation, and matrix quality on forest birds. Ecography 34:103–113. https://doi.org/10.1111/j.1600-0587.2010.06201.x

Steen R (2017) Diel activity, frequency and visit duration of pollinators in focal plants: in situ automatic camera monitoring and data processing. Methods in Ecology and Evolution 8:203–213. https://doi.org/10.1111/2041-210X.12654

Steffan-Dewenter I, Münzenberg U, Bürger C, Thies C, Tscharntke T (2002) Scale□dependent effects of landscape context on three pollinator guilds. Ecology 83:1421–1432. https://doi.org/10.1890/0012-9658(2002)083[1421:SDEOLC]2.0.CO;2

Stephens J, Molan PC, Clarkson BD (2005) A review of Leptospermum scoparium (Myrtaceae) in New Zealand. New Zealand Journal of Botany 43:431–449. https://doi.org/10.1080/0028825X.2005.9512966

Stone GN, Raine NE, Prescott M, Willmer PG (2003) Pollination ecology of acacias (Fabaceae, Mimosoideae). Australian Systematic Botany 16:103–118. https://doi.org/10.1071/SB02024

Thessen A (2016) Adoption of machine learning techniques in ecology and earth science. One Ecosystem 1:e8621. https://doi.org/10.3897/oneeco.1.e8621

Thomas D (1980) Foraging of honeyeaters in an area of Tasmanian sclerophyll forest. Emu 80:55–58. https://doi.org/10.1071/MU9800055

Vanbergen AJ, Insect Pollinators Initiative (2013) Threats to an ecosystem service: pressures on pollinators. Frontiers in Ecology and the Environment 11:251–259. https://doi.org/10.1890/120126

Willcock S, Martínez-López J, Hooftman DA, Bagstad KJ, Balbi S, Marzo A, Prato C, Sciandrello S, Signorello G, Voigt B (2018) Machine learning for ecosystem services. Ecosystem Services 33:165–174. https://doi.org/10.1016/j.ecoser.2018.04.004

Wiltshire R, Potts B (2007) EucaFlip-Life-size guide to the eucalypts of Tasmania. University of Tasmania,

Wimmer J, Towsey M, Roe P, Williamson I (2013) Sampling environmental acoustic recordings to determine bird species richness. Ecological Applications 23:1419–1428. https://doi.org/10.1890/12-2088.1

Winfree R, Aguilar R, Vazquez DP, LeBuhn G, Aizen MA (2009) A meta-analysis of bees’ responses to anthropogenic disturbance. Ecology 90:2068–2076. https://doi.org/10.1890/08-1245.1

Winfree R, Bartomeus I, Cariveau DP (2011) Native Pollinators in Anthropogenic Habitats. Annual Review of Ecology, Evolution, and Systematics 42:1–22. https://doi.org/10.1146/annurev-ecolsys-102710-145042

Woinarski J, Cullen J (1984) Distribution of invertebrates on foliage in forests of south□eastern Australia. Australian Journal of Ecology 9:207–232. https://doi.org/10.1111/j.1442-9993.1984.tb01359.x

