## Supplementary material for "Association between land use, land cover, plant genera, and pollinator abundance in mixed-use landscapes": Online Resources

**Online Resource 1.** List of species observed across the different land-use sites.

| Taxa | Lime bay state reserve | Tasman national park (a) | Tasman national park (b) | Plantation (a) | Plantation (b) | Pasture |
| --- | --- | --- | --- | --- | --- | --- |
| **BIRDS (Honeyeaters)** |  |  |  |  |  |  |
| *Melithreptus affinis* | 6 | 7 |  |  |  |  |
| *Melithreptus validirostris* |  | 6 | 7 | 6 |  |  |
| *Nesoptilotis flavicollis* | 31 | 14 | 7 | 4 |  |  |
| *Anthochaera paradoxa* | 19 | 12 | 3 | 5 |  |  |
| *Anthochaera chrysoptera* | 18 | 5 | 2 |  |  |  |
| *Phylidonyris novaehollandiae* | 30 | 1 |  |  |  |  |
| *Phylidonyris pyrrhopterus* | 11 | 20 | 17 | 45 | 18 |  |
| *Acanthorhynchus tenuirostris* |  | 2 |  | 1 |  |  |
| **BEES** |  |  |  |  |  |  |
| **Native bees** |  |  |  |  |  |  |
| *Exoneura* | 54 | 11 | 55 | 5 | 32 | 27 |
| *Lasioglossum* | 9 |  | 6 |  | 5 | 28 |
| Others | 17 | 1 | 14 | 4 | 4 | 12 |
| **Introduced Bees** |  |  |  |  |  |  |
| *Bombus terrestris* | 1 | 2 | 6 | 2 | 3 | 2 |
| *Apis mellifera* | 22 | 22 | 65 | 13 | 30 | 59 |
| **BEETLES** |  |  |  |  |  |  |
| *Tenebrionidae* |  |  | 6 |  | 2 | 5 |
| *Scarabaeidae* | 41 |  | 48 |  |  |  |
| *Buprestidae* | 1 |  | 4 |  |  |  |
| *Oedemeridae* | 46 | 6 | 53 | 26 | 32 |  |
| *Elateridae* |  |  | 1 |  |  |  |
| *Cantharidae* |  |  |  |  | 101 |  |
| *Lycidae* | 13 |  | 17 |  | 2 |  |
| *Cleridae* | 2 |  | 11 |  |  |  |
| *Cerambycidae* | 5 |  | 1 |  |  |  |
| **BUTTERFLIES** |  |  |  |  |  |  |
| *Pieris rapae* | 1 |  |  | 3 | 4 | 14 |
| *Heteronympha merope* |  |  | 16 | 1 | 4 |  |
| *Vanessa itea* |  |  | 3 | 1 | 2 |  |
| *Vanessa kershawi* |  |  | 2 | 2 | 9 |  |
| *Junonia villida* |  |  |  |  |  | 4 |
| *Neolucia agricola* |  |  | 1 |  |  |  |
| *Graphium macleayanum* |  |  |  | 7 |  |  |
| *Hesperilla donnysa* |  |  |  | 10 |  |  |

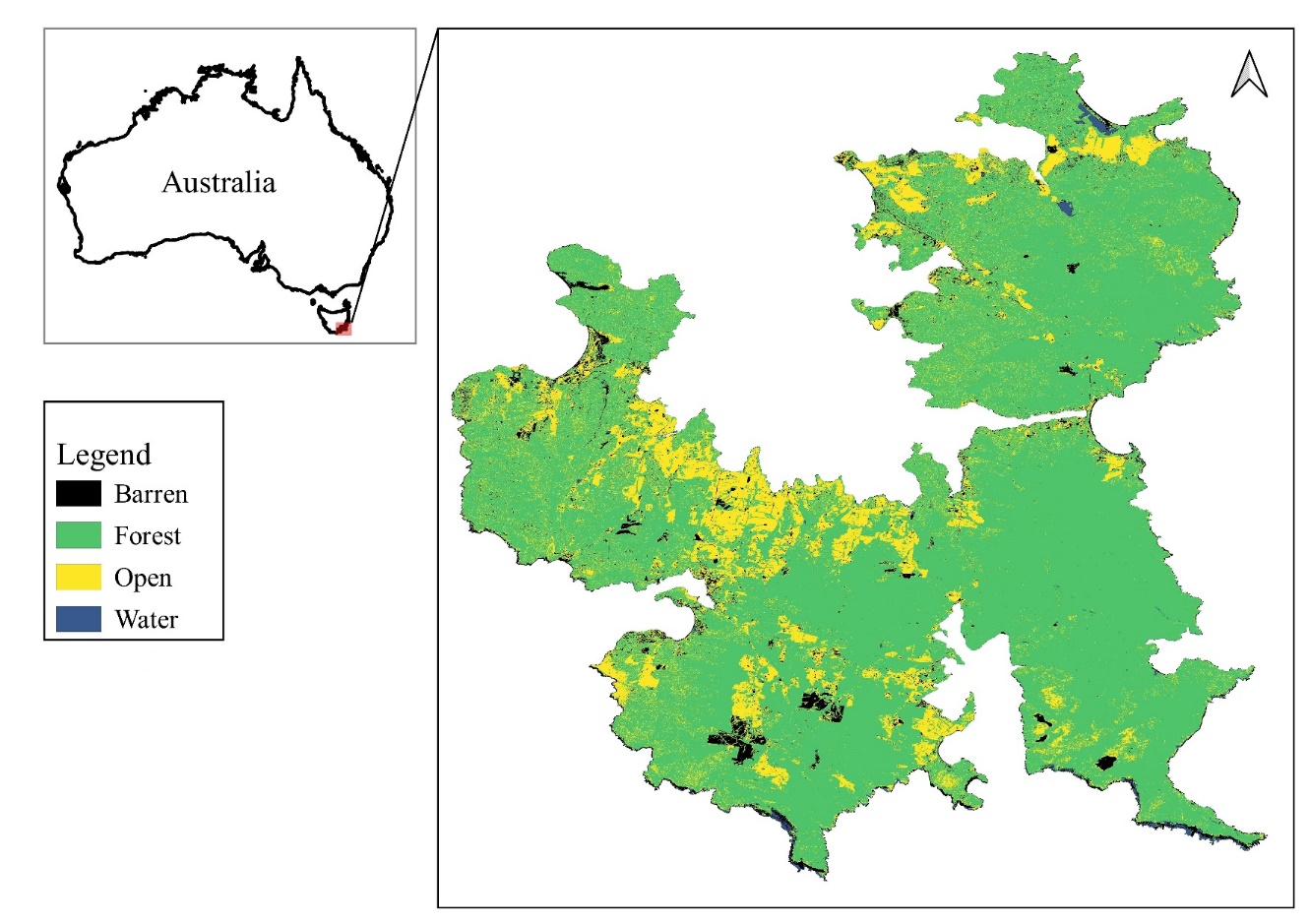

**Online Resource 2.** Land-cover of the Tasman and Forestier Peninsula with forest (green), open (yellow), baren (black) and water (blue) classes.

**Online Resource 3.** Confusion matrix of the classified land-cover image which includes forest, open, baren and water classes. Reference and prediction refer to the validation data and the predicted land-cover image, respectively.

|  | | Reference | | | |
| --- | --- | --- | --- | --- | --- |
|  |  | Forest | Open | Barren | Water |
| Prediction | Forest | 96 | 2 | 2 | 0 |
|  | Open | 6 | 87 | 7 | 0 |
|  | Barren | 1 | 19 | 79 | 1 |
|  | Water | 28 | 16 | 0 | 56 |

**Online Resource 4.** Sensitivity and specificity of each of the land-cover classes

|  | Forest | Open | Barren | Water |
| --- | --- | --- | --- | --- |
| Sensitivity | 0.7328 | 0.7016 | 0.8977 | 0.9825 |
| Specificity | 0.9851 | 0.9529 | 0.9327 | 0.8717 |

**Online Resource 5.** Percentage of forest and open land cover at 100 m and 2000 m radius from the centre of the 2-ha plot. Percentage of barren and water later cover not included.

| Plot code | Forest.100 | Open.100 | Forest.2000 | Open.2000 |
| --- | --- | --- | --- | --- |
| EX2 | 99.684 | 0.316 | 84.655 | 6.992 |
| EX3 | 100.000 | 0.000 | 90.661 | 5.508 |
| EX4 | 100.000 | 0.000 | 92.159 | 5.930 |
| EY1 | 98.722 | 0.000 | 89.839 | 6.746 |
| EY2 | 100.000 | 0.000 | 87.116 | 10.204 |
| EY3 | 98.403 | 0.639 | 92.664 | 6.252 |
| FX0 | 98.071 | 1.929 | 96.430 | 1.936 |
| FX2 | 99.359 | 0.641 | 96.966 | 1.145 |
| FX4 | 100.000 | 0.000 | 96.161 | 1.570 |
| FX7 | 95.833 | 3.205 | 95.804 | 1.347 |
| FX8 | 90.260 | 5.195 | 96.815 | 1.939 |
| FX9 | 89.068 | 7.717 | 96.461 | 2.390 |
| LX1 | 92.698 | 7.302 | 82.053 | 12.182 |
| LX2 | 91.667 | 8.013 | 81.363 | 11.759 |
| LX3 | 88.535 | 11.465 | 79.081 | 12.504 |
| LY1 | 93.910 | 6.090 | 84.440 | 12.910 |
| LY2 | 94.340 | 5.660 | 86.445 | 11.730 |
| LY4 | 82.595 | 17.405 | 86.552 | 11.493 |
| NX1 | 98.397 | 1.603 | 68.320 | 18.953 |
| NX2 | 96.190 | 3.175 | 69.121 | 17.524 |
| NX3 | 97.785 | 2.215 | 66.726 | 18.851 |
| NY1 | 84.444 | 15.556 | 71.584 | 16.345 |
| NY2 | 47.588 | 11.254 | 68.299 | 17.359 |
| NY3 | 59.683 | 15.556 | 69.103 | 17.427 |
| PX2 | 3.165 | 96.835 | 42.214 | 54.631 |
| PX3 | 20.755 | 75.157 | 46.953 | 50.194 |
| PX4 | 6.731 | 93.269 | 52.135 | 45.565 |
| PY2 | 10.759 | 79.747 | 36.272 | 59.115 |
| PY3 | 0.000 | 91.429 | 37.133 | 58.661 |
| PY4 | 6.013 | 84.494 | 39.258 | 57.082 |
| TX1 | 77.636 | 20.447 | 80.613 | 15.150 |
| TX2 | 84.227 | 15.457 | 82.653 | 14.782 |
| TX3 | 99.045 | 0.955 | 81.066 | 17.461 |
| TY0 | 93.930 | 6.070 | 71.601 | 25.976 |
| TY1 | 88.782 | 11.218 | 74.124 | 23.885 |
| TY2 | 94.586 | 5.414 | 81.080 | 17.909 |

**Online Resource 6.** Percentage of forest and open land cover at 500 m and 2000 m radius from the centre of the 1000 m transect. Percentage of barren and water later cover not included.

| Plot Code | Forest.500 | Open.500 | Forest.2000 | Open.2000 |
| --- | --- | --- | --- | --- |
| LX | 77.249 | 9.912 | 81.024 | 12.027 |
| LY | 86.959 | 12.523 | 85.824 | 12.359 |
| PY | 22.144 | 73.394 | 37.189 | 58.645 |
| PX | 38.653 | 58.661 | 43.592 | 53.299 |
| TY | 83.142 | 15.940 | 73.906 | 24.118 |
| TX | 92.745 | 7.038 | 82.734 | 14.501 |
| NX | 90.460 | 8.929 | 68.714 | 17.805 |
| NY | 46.654 | 6.858 | 68.370 | 17.282 |
| EX | 99.491 | 0.395 | 90.363 | 5.447 |
| EY | 94.369 | 0.599 | 92.727 | 4.784 |

**Online Resource 7.** The RMSE and R2 and RMSE of the different butterfly count model along with the variable importance (gini values) of the different predictors.

| **Model** | **Predictors** | **RMSE** | **R2** | **Predictors** | **Gini values** |
| --- | --- | --- | --- | --- | --- |
| Butterfly count model | pasture + plantation+ protected.area + forest.500 + open.500 | 3.448 | 0.019 | forest.500 | 121.730 |
|  |  |  |  | open.500 | 96.567 |
|  |  |  |  | plantation | 13.468 |
|  |  |  |  | protected.area | 12.104 |
|  |  |  |  | pasture | 3.441 |

**Online Resource 8.** Variable importance (gini values) of the top 10 predictors of the different pollinator count models

| **Model** | **Predictors** | **Gini values** |
| --- | --- | --- |
| Honeyeater count | 1. forest.2000 | 136.492699 |
|  | 2. open.2000 | 134.955890 |
|  | 3. Eucalyptus | 91.420123 |
|  | 4. Symphyomyrtus | 30.876710 |
|  | 5. LU_protected.area | 24.544626 |
|  | 6. LU_pasture | 19.752531 |
|  | 7. LU_plantation | 5.531838 |
| Native bee count | 1. forest.2000 | 59.91558522 |
|  | 2. genera_Pultenaea | 57.62292383 |
|  | 3. open.2000 | 54.30421052 |
|  | 4. genera_Leucopogon | 53.99463337 |
|  | 5. genera_Acacia | 18.81130382 |
|  | 6. genera_Goodenia | 10.03147224 |
|  | 7. genera_Pimelea | 7.80471331 |
|  | 8. genera_Olearia | 6.68817358 |
|  | 9. LU_pasture | 5.96508320 |
|  | 10. genera_Leptospermum | 5.11976439 |
| Introduced bee count | 1. open.100 | 38.05639442 |
|  | 2. forest.100 | 36.27041769 |
|  | 3. genera_Acacia | 19.88738334 |
|  | 4. genera_Melaleuca | 18.74869533 |
|  | 5. genera_Pomaderris | 12.88385697 |
|  | 6. genera_Pimelea | 11.96128822 |
|  | 7. genera_Lissanthe | 6.00771167 |
|  | 8. genera_Pultenaea | 5.55470149 |
|  | 9. LU_protected.area | 3.59441529 |
|  | 10. LU_plantation | 3.28012755 |
| Beetle count | 1. genera_Leptospermum | 2.162004e+03 |
|  | 2. forest.100 | 7.262206e+02 |
|  | 3. open.100 | 6.908810e+02 |
|  | 4. LU_plantation | 1.155066e+02 |
|  | 5. LU_protected.area | 9.003934e+01 |
|  | 6. genera_Pultenaea | 7.575783e+01 |
|  | 7. genera_Pomaderris | 4.617187e+01 |
|  | 8. genera_Acacia | 3.255777e+01 |
|  | 9. genera_Pimelea | 2.400601e+01 |
|  | 10. LU_pasture | 2.212450e+01 |
